# Mendelian inheritance errors in whole genome sequenced trios are enriched in repeats and cluster within copy number losses

**DOI:** 10.1101/240424

**Authors:** Prachi Kothiyal, Wendy S. Wong, Dale L. Bodian, John E. Niederhuber

**Author notes:** Corresponding authors; Prachi Kothiyal John E. Niederhuber.

## Abstract

Trio-based whole genome sequencing (WGS) data can contribute significantly towards the development of quality control methods that can be applied to non-family WGS. Mendelian inheritance errors (MIEs) in parent-offspring trios are commonly attributed to erroneous sequencing calls, as the rate of true de novo mutations is extremely low compared to the incidence of MIEs. Here, we analyzed WGS data from 1,314 trios across diverse human populations with the goal of studying the characteristics of MIEs. We applied filters based on genotype call quality and observed that filtering has a greater impact on frequent MIEs. Our results indicate that MIEs are enriched in repeats and MIE density correlates with short interspersed nuclear elements (SINEs) density. We also observed clustered MIEs in regions overlapping large deletions. We created population-specific MIE profiles and discovered regions that represent different MIE distributions across populations. Finally, we have provided population-specific MIE tracks that can be loaded in UCSC Genome Browser. These profiles can be used for flagging calls in proximity of clustered MIEs before allele frequency and admixture calculations, annotating candidate de novo mutations, discovering population-specific putative deletions, and for distinguishing between regions that have errors due to sequence quality vs. chromosomal anomalies.

## Introduction

Next-generation sequencing (NGS) technologies are increasingly used for the high-throughput detection of variants in the genome (Bentley et al. 2008) and are enabling advances in the clinical application of genomics. However, NGS technologies are often associated with a non-negligible rate of incorrectly identified sequence variants which are a result of sequencing errors instead of true genetic variation (Jünemann et al. 2013; Loman et al. 2012; Dewey et al. 2014; Ross et al. 2013; O’Rawe et al. 2013). Previous findings have suggested an error rate of ⍰00.1−1 × 10−2 per base sequenced using an Illumina platform (Loman et al. 2012; Meacham et al. 2011). An error rate of this magnitude translates to millions of incorrect calls per human genome sequenced. Previous research studies have explored various mechanisms of mitigating these errors. Approaches range from optimizing library preparation techniques to deploying downstream in silico filters (Lou et al. 2013; Reumers et al. 2011).

A Mendelian inheritance error (MIE) represents a genotype in the offspring that could not have been inherited from either biological parent by Mendelian inheritance with the assumed ploidy. MIEs can arise from de novo mutations that could be germline or non-germline, genotype calling errors in the child or parent, or incorrect pedigree information. The de novo mutation rate for humans has been estimated to be ~1.2e-8 per generation, which translates to an estimated 38 mutations per offspring (Kong et al. 2012; Goldmann et al. 2016; Wong et al. 2016; Conrad et al. 2011). With rate of true de novo mutations being around 4 orders of magnitude lower than MIE rate in our cohort (Goldmann et al. 2017), most Mendelian inconsistencies can be attributed to sequencing errors or sequence variation due to chromosomal anomalies that manifest as Mendelian inconsistent calls.

While MIEs have been evaluated before (Patel et al. 2014; Pilipenko et al. 2014; Blue et al. 2014), these studies are performed on a smaller scale in terms of number of trios and coverage across genomes and populations. Existing tools can be used to reduce false positive calls in family-based sequencing data by checking genotyping calls for consistency (Abecasis et al. 2002; Douglas et al. 2000; Sobel et al. 2002; O’Connell and Weeks 1998). However, most approaches are designed with the primary goal of discarding inconsistent calls as sequencing errors instead of exploring their properties as indicators of larger variations. While SNP genotyping data from parent-offspring trios has been used for detection of deletion polymorphisms (Conrad et al. 2006), a similar approach has not been explored extensively for WGS trio data (Manheimer et al. 2017).

We aim to bridge the gap between MIE detection and their removal or adoption in alternative applications by first understanding their characteristics. Using a trio design for 1,314 families, we present an overview of the distribution and characteristics of MIEs. We explore whether comprehensive analysis of MIEs can contribute to the development of filtering approaches for mitigating sequencing artifacts. We evaluate call quality metrics such as read depth, genotype quality, and allelic balance to assess their impact on the rate of MIEs. Finally, we demonstrate that MIE profiles in the autosomes, especially regions enriched for errors unique to a trio or a population, can be used to discover large deletions.

With the implication of de novo mutations in disorders ranging from rare congenital syndromes to relatively common disorders such as epilepsy (Epi4K Consortium et al. 2013), autism (Ronemus et al. 2014; Yuen et al. 2016), severe intellectual disability (Rauch et al. 2012), and late-onset phenotypes such as schizophrenia (McCarthy et al. 2014), they are being studied more routinely. We provide population-specific MIE tracks that can be used for annotating MIEs with frequencies in a larger population to aid in the selection of candidate de novo mutations.

## Results

### Filters based on variant call quality have different impact on rare vs. frequent MIEs

We generated MIE map for each autosome by including all variant sites where an MIE is observed in at least one trio. The terminology used for describing overall statistics is explained in Figure 1a. We observed ~3.29 million unique sites across the autosomes containing at least one Mendelian inconsistent genotype, with ~0.78 million sites where the Mendelian violation exists in only one trio (Table 1). On average, 56 out of 1,314 trios were found to have an inconsistent genotype at a MIE site.

**Table 1:** Overall MIE statistics before and after applying quality-based filters

|  | **Pre-filtering**<br>(95% confidence<br>interval where<br>applicable) | **Post-filtering**<br>(95% confidence<br>interval where<br>applicable) |
| --- | --- | --- |
| **Cohort summary statistics** |  |  |
| Total no. of errors | 214,415,087 | 29,857,521 |
| Total no. of sites | 3,829,459 | 1,864,004 |
| Total no. of sites with 1<br>inconsistent trio | 783,673 | 595,218 |
| Average no. of inconsistent<br>trios per MIE site | 56<br>(50-62) | 16<br>(14-18) |
| **Trio summary statistics** |  |  |
| Average % variant sites<br>with MIE | 4.56<br>(4.55-4.57) | 0.63<br>(0.626-0.633) |
| Average % unique MIEs in<br>each trio | 0.36<br>(0.35-0.37) | 1.97<br>(1.92-2.02) |
| Average no. of MIEs per trio | 162,929<br>(162,539-163,318) | 22,673<br>(22,539-22,806) |
| Average no. of MIEs unique<br>to trio | 595<br>(576-614) | 453<br>(440-446) |

Given that on average ~4.5% variant sites in a trio contained an MIE, we aimed to determine whether the Mendelian inconsistencies are likely caused by sequencing error. We used previously determined cutoffs for Genotype Quality (GQ), Read Depth (DP), and Allele Balance (AB) at 30, 10, and 0.25 (Bodian et al. 2016) respectively to determine the impact of quality-based filtering on MIE distribution. Filtering variants by quality metrics reduced the total number of MIEs by ~86% (Table 1) while reducing the number of unique errors (sites with MIE in a single trio) by ~24%. On average, 0.63% variant sites contained MIEs per trio after filtering. Filtering impacts a larger proportion of higher frequency MIEs when compared to unique MIEs (Supplementary Figure 1). We hypothesized that errors resulting from systematic issues (such as paralogs) would be found in a higher number of genomes, whereas unique errors could result from family-specific variation, either de novo mutations or larger inherited events (Conrad et al. 2006; Weir 2012), or from misincorporation of a base. These results suggest that filtering the genotypes by quality metrics may distinguish MIEs that result from sequencing error from those that may be capturing true sequence variation.

While quality-based filtering reduced the number of Mendelian-inconsistent calls, we also assessed distribution of quality metrics across Mendelian-consistent calls. For each autosomal position with an MIE in at least one trio, we extracted Mendelian-consistent and –inconsistent genotypes (Figure 1a) and corresponding GQ, DP, and AB across the entire cohort. The data from chromosome 16 are plotted in Figure 1b as an example. We evaluated minimum GQ, DP, and AB (only for heterozygous calls) values across trio members instead of including all three values or using averages because an erroneous call in any of the three members can lead to a MIE and therefore genotype with lowest quality has a higher likelihood of being the culprit. Mendelian-inconsistent trios have a higher proportion of calls at low GQ and DP compared to consistent trios (Figure 1b). However, we observed peaks at high AB for Mendelian inconsistent calls. To determine if this was caused by calls with poor quality where sufficient evidence was not available to make a heterozygous call, we generated the distribution for minimum AB in trios after filtering calls on GQ≥30 and DP≥10. This led to a reduction in the proportion of MIEs with high AB (Supplementary Figure 2) implying that heterozygous calls with AB close to 1 tend to be associated with low call quality.

**Figure 1a:**
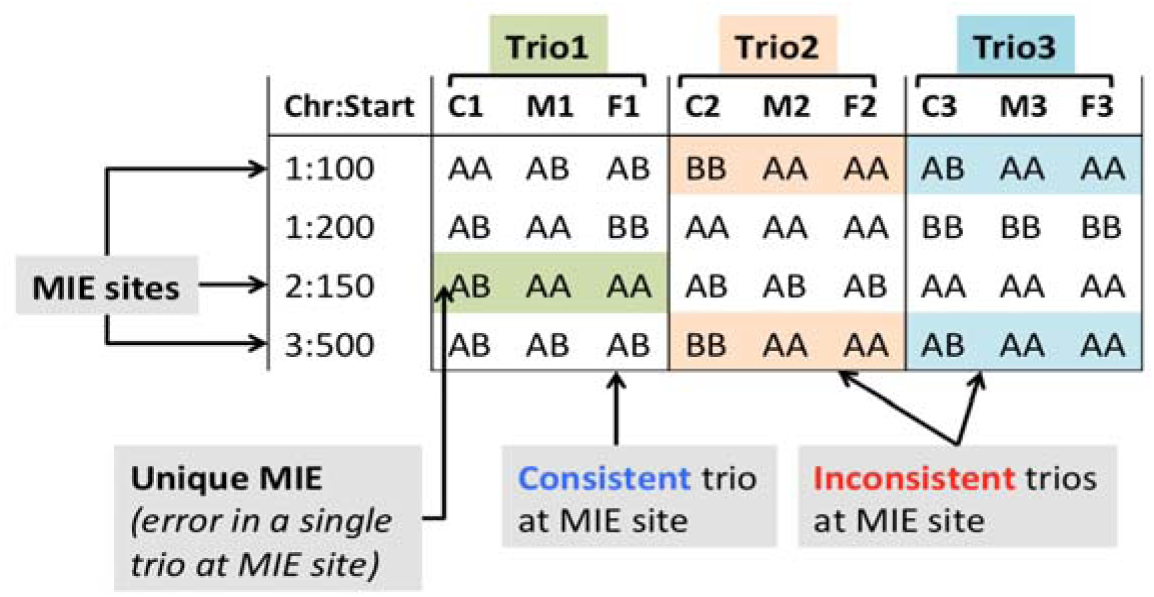
Example illustrating term definitions for MIEs. A locus that has at least one trio with Mendelian inconsistent genotype calls is defined as an “MIE site”. A “unique MIE” occurs when at any given MIE site only a single trio has inconsistent calls. “Mendelian-inconsistent trios” and “Mendelian-consistent trios” contain inconsistent and consistent calls respectively at a given MIE site.

**Figure 1b:**
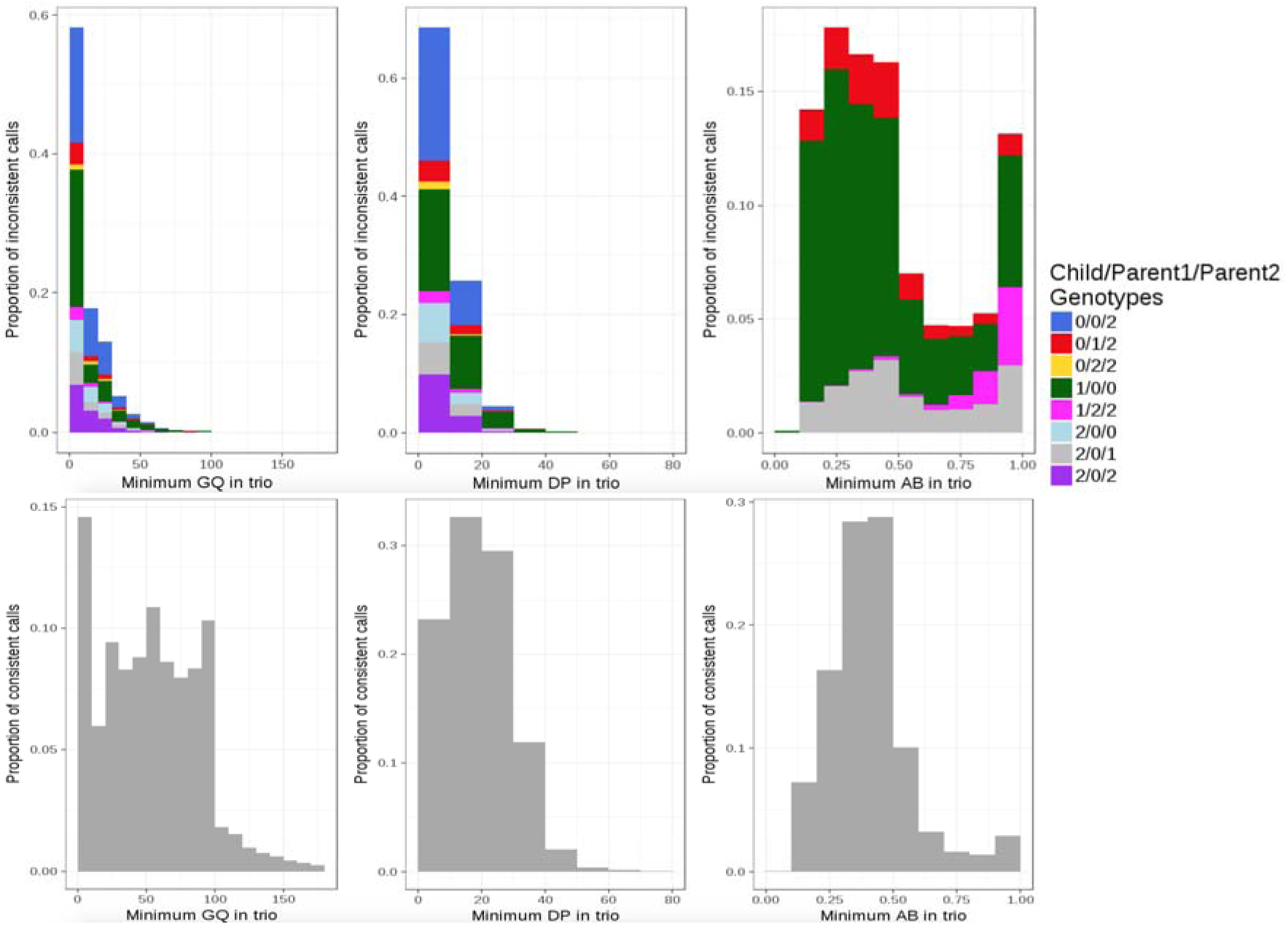
Distribution of call quality metrics for MIE sites in chromosome 16. The top row contains histograms for minimum GQ (within a given trio), minimum DP, and minimum AB (across heterozygous calls in trio) in Mendelian-inconsistent trios for chromosome 16 MIE sites. X-axis is the quality metric being considered and Y-axis is the proportion of total MIEs across all sites and trios that correspond to the bin on X-axis. The bars are filled with colors depicted in the legend corresponding to the combination of genotype calls in child and parents where “0”, “1”, and “2” map to AA, AB, and BB genotypes respectively. The genotype classification does not distinguish between maternal and paternal genotypes (e.g., Child/Mother/Father genotypes AA/AB/BB and AA/BB/AB are combined). The bottom row illustrates distributions for Mendelian-consistent trios at MIE sites.

### MIE counts across autosomes and enrichment in repeat regions

Since one possible source of MIEs is repeat sequences (Pilipenko et al. 2014; Treangen and Salzberg 2012), we tested whether the MIEs lie in regions enriched in repetitive sequence or repeat elements. An average of 72.84% (95% CI: 72.6%, 73%) of the total MIE sites across each autosome overlap with repeat elements (Table 2 and Supplementary Figure 3). Chromosome 19 has the highest MIE density at 2.25 sites per kb of its length and also has the highest number of MIEs per 1000 variant sites, at 36.43.

**Table 2:** Overall MIE statistics for each autosome

| Autosome | Variants<br>(x10<sup>6</sup>) | MIE<br>sites | No. of MIE<br>sites per<br>1000<br>variant<br>sites | MIE sites<br>overlapping<br>repeats | % MIE sites<br>overlapping<br>repeats | No. of MIE<br>sites per 1Kb |
| --- | --- | --- | --- | --- | --- | --- |
| chr1 | 13.21 | 305810 | 23.15 | 225039 | 73.59 | 1.23 |
| chr2 | 13.51 | 290446 | 21.50 | 213961 | 73.67 | 1.19 |
| chr3 | 10.29 | 231046 | 22.45 | 179292 | 77.60 | 1.17 |
| chr4 | 10.48 | 258814 | 24.70 | 199532 | 77.09 | 1.35 |
| chr5 | 10.39 | 217360 | 20.92 | 166580 | 76.64 | 1.20 |
| chr6 | 9.05 | 235084 | 25.97 | 177011 | 75.30 | 1.37 |
| chr7 | 10.08 | 237449 | 23.56 | 172838 | 72.79 | 1.49 |
| chr8 | 8.75 | 201867 | 23.07 | 144497 | 71.58 | 1.38 |
| chr9 | 10.71 | 233368 | 21.79 | 161262 | 69.10 | 1.65 |
| chr10 | 8.42 | 188857 | 22.44 | 135697 | 71.85 | 1.39 |
| chr11 | 7.50 | 180554 | 24.08 | 138832 | 76.89 | 1.34 |
| chr12 | 6.89 | 170713 | 24.77 | 131946 | 77.29 | 1.28 |
| chr13 | 4.77 | 117183 | 24.56 | 85392 | 72.87 | 1.02 |
| chr14 | 5.08 | 129636 | 25.54 | 93466 | 72.10 | 1.21 |
| chr15 | 6.23 | 141241 | 22.68 | 94671 | 67.03 | 1.38 |
| chr16 | 6.09 | 137433 | 22.56 | 95424 | 69.43 | 1.52 |
| chr17 | 4.62 | 124768 | 27.03 | 88445 | 70.89 | 1.54 |
| chr18 | 3.78 | 87674 | 23.22 | 64659 | 73.75 | 1.12 |
| chr19 | 3.66 | 133308 | 36.43 | 99930 | 74.96 | 2.25 |
| chr20 | 3.03 | 75515 | 24.89 | 55728 | 73.80 | 1.20 |
| chr21 | 1.88 | 54204 | 28.80 | 38221 | 70.51 | 1.13 |
| chr22 | 2.67 | 77129 | 28.90 | 49101 | 63.66 | 1.50 |

To further characterize the relationship between MIEs and repeats, we compared the positions of the MIEs to different types of repeat elements. Long interspersed nuclear elements (LINEs) have the highest genomic coverage (21%) followed by short interspersed nuclear elements (SINEs) (15%) (Treangen and Salzberg 2012). However, SINEs contained the highest percentage of filtered MIE sites (35%), with LINEs being second highest at 21%. Chromosome 19 has been studied previously for its highest repeat density (nearly 55%) among all autosomes and also its high content of SINEs (Grimwood et al. 2004). We assessed the relationship between different repeat types and MIE density. Linear regression of SINE density against MIE density produced an R-squared of 0.67 with a p-value <2.2e-16 (Figure 2). Excluding chromosome 19 resulted in a lower R-squared (0.4, p-value <2.2e-16) but the correlation was still stronger when compared to other repeat types (Supplementary Table 1). In contrast, we obtained an R-squared of 0.16 with linear regression for MIE density against cumulative repeat density including all repeat types.

**Figure 2:**
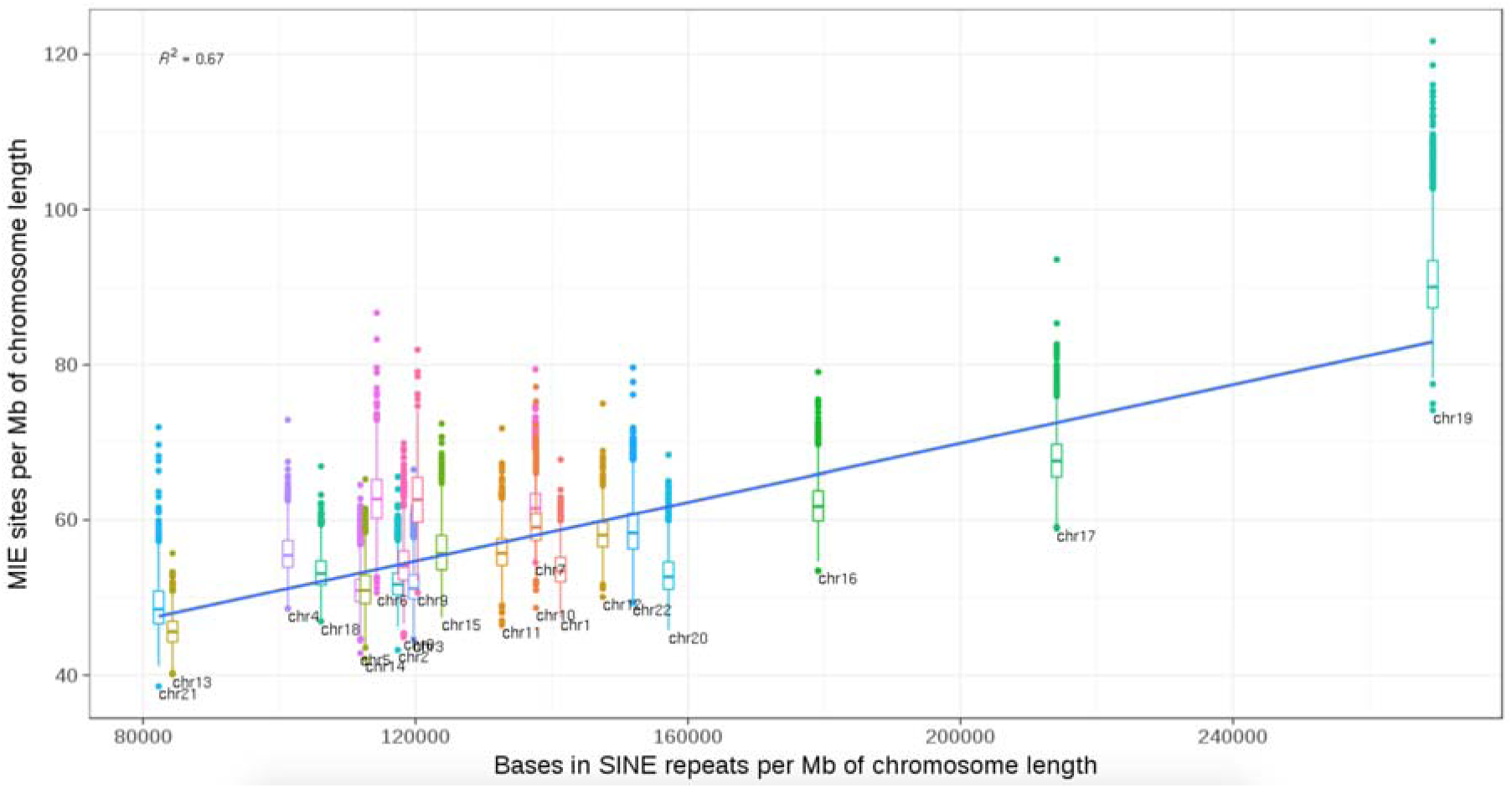
MIE density (number of MIEs per Mb of autosome) against SINE density for each autosome across all trios. Linear regression line for SINE density (X-axis) against MIE density (Y-axis) is in blue. Each boxplot represents data points for a single autosome and each point represents a single trio.

As SINEs are primarily comprised of Alu elements, we evaluated the contribution of different Alu types (sub-families AluJ, AluS, and AluY, ordered from most ancient to youngest) to MIE counts pre- and post-filtering (Supplementary Figure 4). AluS is the most abundant sub-family covering ~6.4% of the genome (Grover et al. 2004) and ~19% of total MIE sites overlap AluS elements. AluJ and AluY contained 5% and 7.25% respectively of the total MIE sites. Due to their structure and abundance (Lander et al. 2001; Deininger 2011), Alus are difficult to sequence, align and call variants in, and therefore variants within Alus are usually filtered out. However, further work is needed to understand if MIEs within Alus can be used to detect deletion or insertion of polymorphic Alus with respect to the reference genome (Hormozdiari et al. 2011).

### MIE enrichment correlates with copy number losses

High frequency MIEs could indicate systematic errors in the technology or common copy number variations not represented in the reference. However, MIEs that are observed at a low frequency could be due to less frequent copy number variation or chromosomal anomalies and the underlying variant calls may not always be associated with poor quality (Weir 2012; Conrad et al. 2006). An example of MIE due to a heterozygous deletion in a parent and child is illustrated in Figure 3a. In such cases, MIEs can be indicative of an underlying anomaly instead of being sequencing artifacts.

**Figure 3a:**
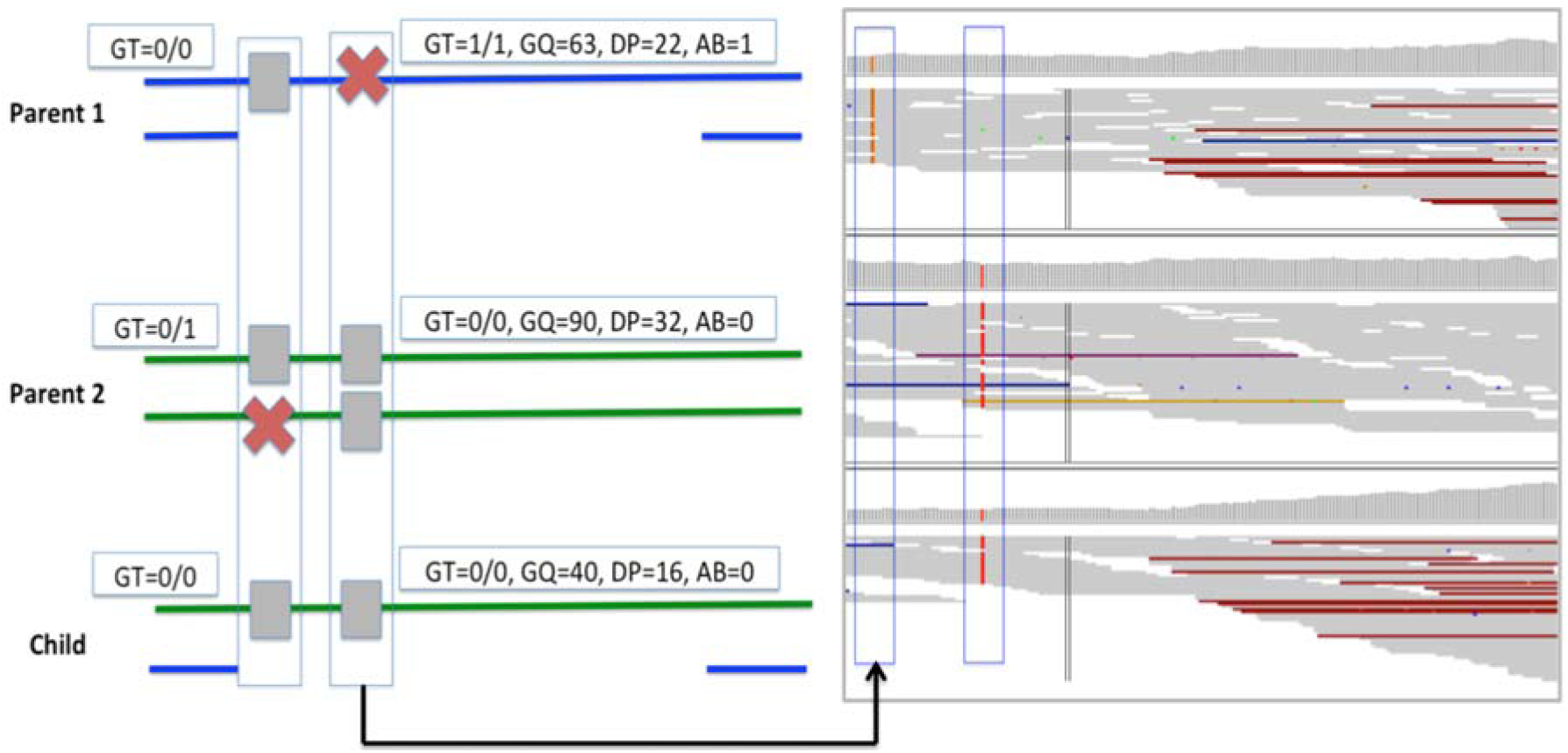
A real example of MIE call that overlaps a large deletion and passes quality-based filters. Left panel depicts an MIE call in the trio along with quality metrics for the genotype call in each member of the trio. Parent 1 has a deletion in the bottom allele that is also inherited by the child. Parent 2 does not have the deletion. Panel on the right presents a screenshot of IGV visualization of the alignment for the site and surrounding context. Two MIE calls in the trio are highlighted within blue rectangles where the first box corresponds to the MIE signature in the left panel. A Mendelian consistent call overlapping the deletion is also illustrated in the figure where Parent 1 and Child are called homozygous reference instead of hemizygous.

We utilized standardized scores for unique and total MIEs to filter trios that have a disproportionately high number of unique MIE sites in any given autosome (Figure 3b). We selected the trios with highest scores for unique and total MIEs and assessed if these trios had clusters of unique MIEs in any 1Mb window along the chromosome length (Supplementary Table 2). An example is shown for chromosome 10 in Supplementary Figure 5 where top panel shows the distribution of unique MIE calls and bottom panel shows the breakpoints for corresponding copy number loss. We also compared the calls against results obtained with Manta (Chen et al. 2016) and Control-FREEC (Boeva et al. 2012). Of the 22 copy number losses, 14 were detected with both tools, 6 were detected only with Freec, 1 was detected only with Manta and 1 was detected only with presence of clustered MIEs. Based on alignment data and the presence of potential breakpoints, these cases could be considered as candidates for further validation to confirm the presence of copy number losses.

**Figure 3b:**
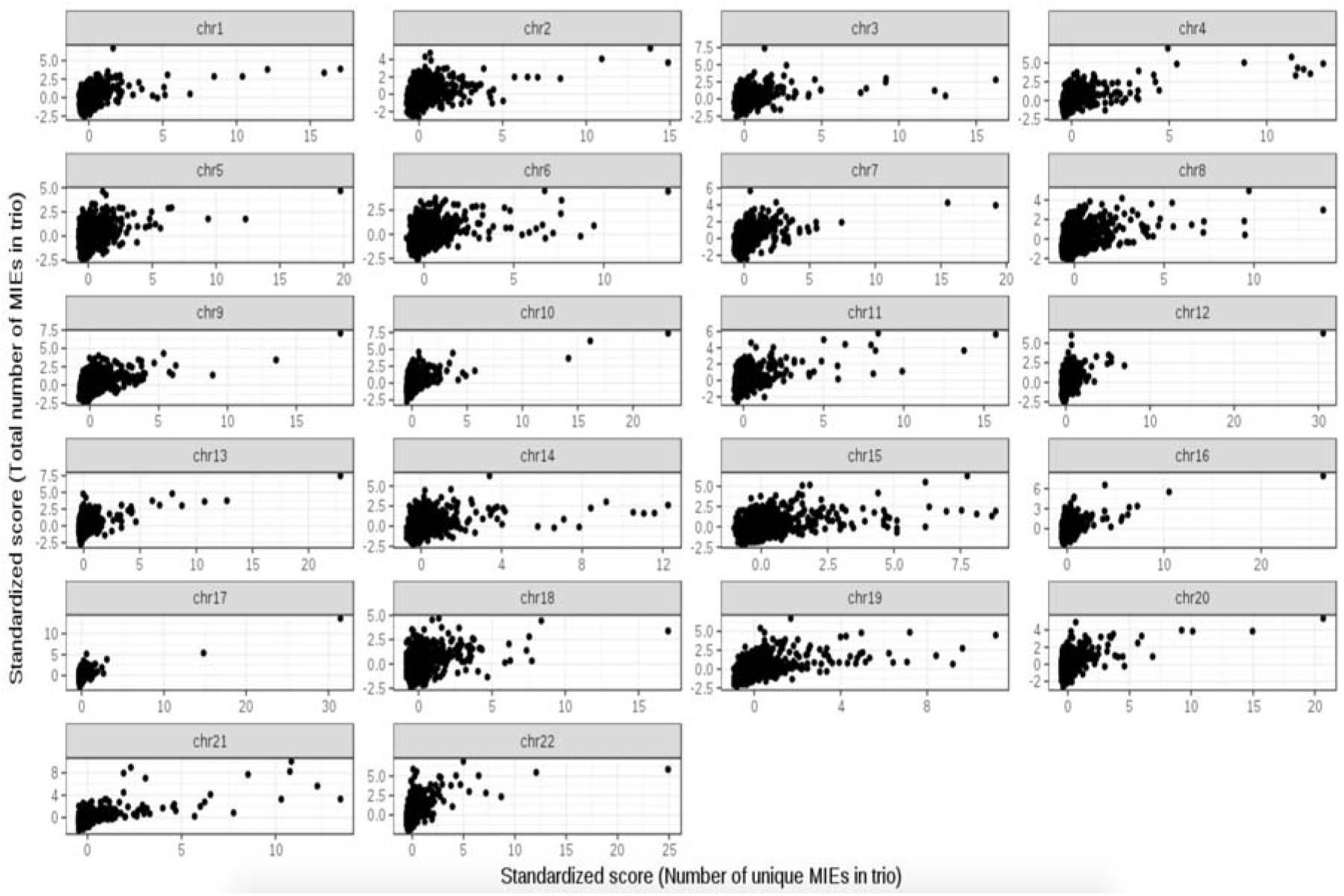
Distribution of Z-scores for total number of MIE sites vs. Z-scores for number of unique MIE sites for all trios in each autosome. Each data point represents a single trio. Standardized scores for MIE counts are on the Y-axis and Standardized score for number of unique MIE sites on the X-axis per autosome is on the X-axis.

## Population-specific MIE profiles

We compared MIE distribution against NIST high confidence regions (HCR) (Zook et al. 2014) and 1000Genomes Structural Variation (SV) map (Sudmant et al. 2015) to assess if regions with MIEs are excluded from NIST HCR (Table 3) or if MIEs are enriched in regions overlapping known deletions. We have used Chromosome 16 as an example to describe the characteristics of MIEs in our cohort (Figure 4a). While some of the MIE peaks correspond to regions excluded from NIST HCR, some are included. In order to determine if MIE-enriched regions in ITMI data are included in NIST HCR due to population-specific differences, we separated MIE calls by ethnicity. We observed population-specific peaks in MIE distributions both before and after filtering. An example is chr16:6.7MB-6.9MB where profile for AMR trios showed a higher MIE count compared to EUR trios and was also annotated as a deletion in 1000Genomes SV map (highlighted in red in Figure 4a). We aimed to determine if the higher number of MIE sites in AMR population could be attributed to a subset of trios or if all trios showed similar enrichment. Three AMR trios contained a high number of unique MIE sites as well as total number of MIEs in chr16:6.7MB-6.9MB when compared to other populations (Supplementary Figure 6a). In contrast, a peak common to all populations and excluded from NIST HCR (chr16:55.7MB-55.9MB; highlighted in blue in Figure 4a) was associated with a higher number of MIE sites across all populations but did not have outliers due to high number of unique sites (Supplementary Figure 4a). Child and father in all 3 trios with high number of unique MIEs in chr16:6.7MB-6.9MB contain a ~180Kb heterozygous deletion in the region (Supplementary Figure 6b) and we could also confirm this deletion against calls generated with Manta.

**Figure 4a:**
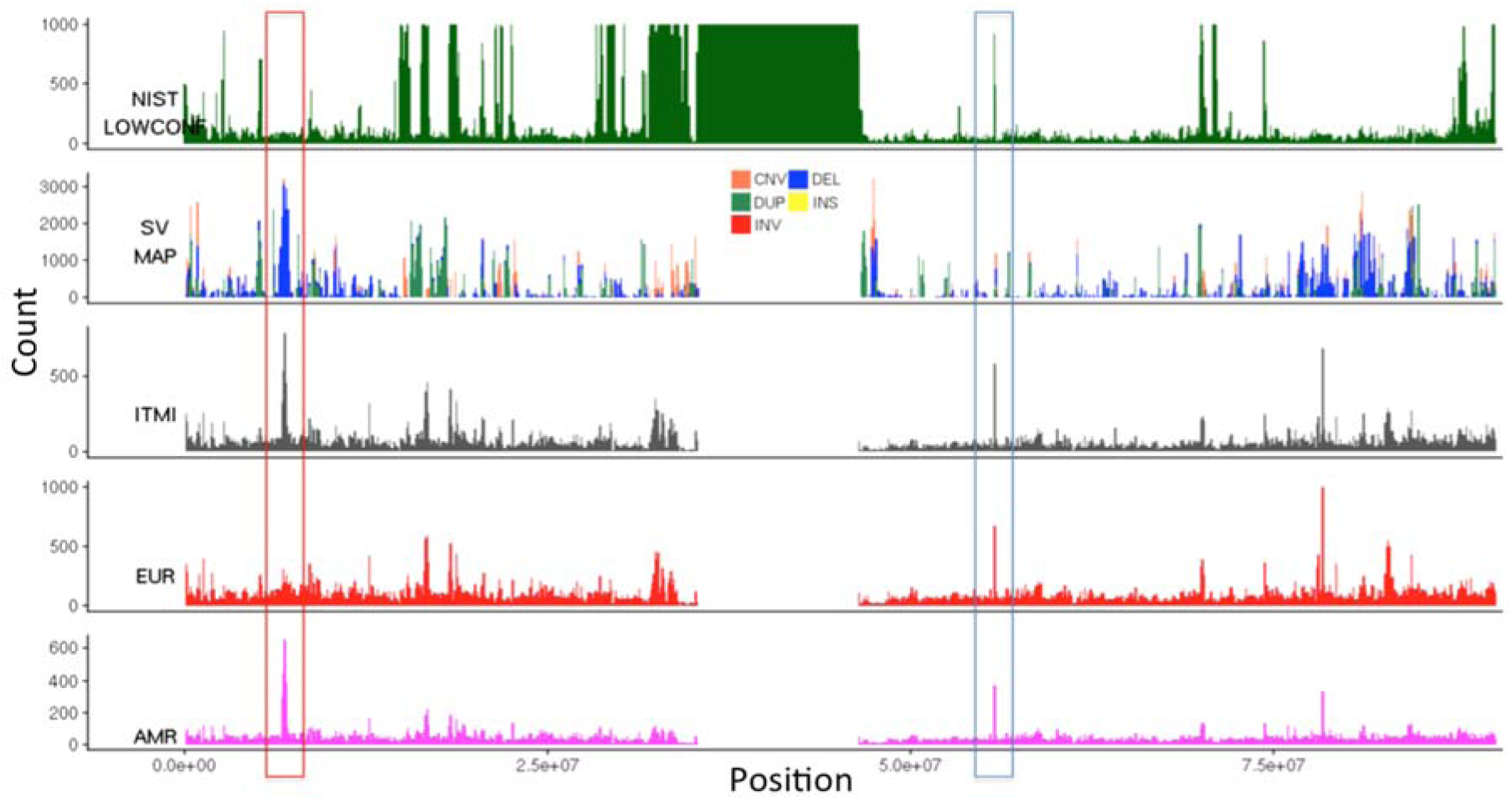
Tracks for NIST low confidence regions (top), 1000Genomes SVs (2^nd^ from top), MIEs in ITMI data post-filtering, and MIEs in EUR and AMR populations for Chromosome 16. Height of a peak represents how many sites in the 100Kb window are excluded from NIST HCR, overlap with SVs, or contain MIEs in ITMI data.

We created bedGraph for each autosome and population to summarize MIE sites and associated error rates specific to the population. These are provided as Supplementary Files in an accompanying zipped archive. The bed files can be loaded as tracks and viewed in UCSC Genome Browser (Kent et al. 2002) (Figure 4b).

**Figure 4b:**
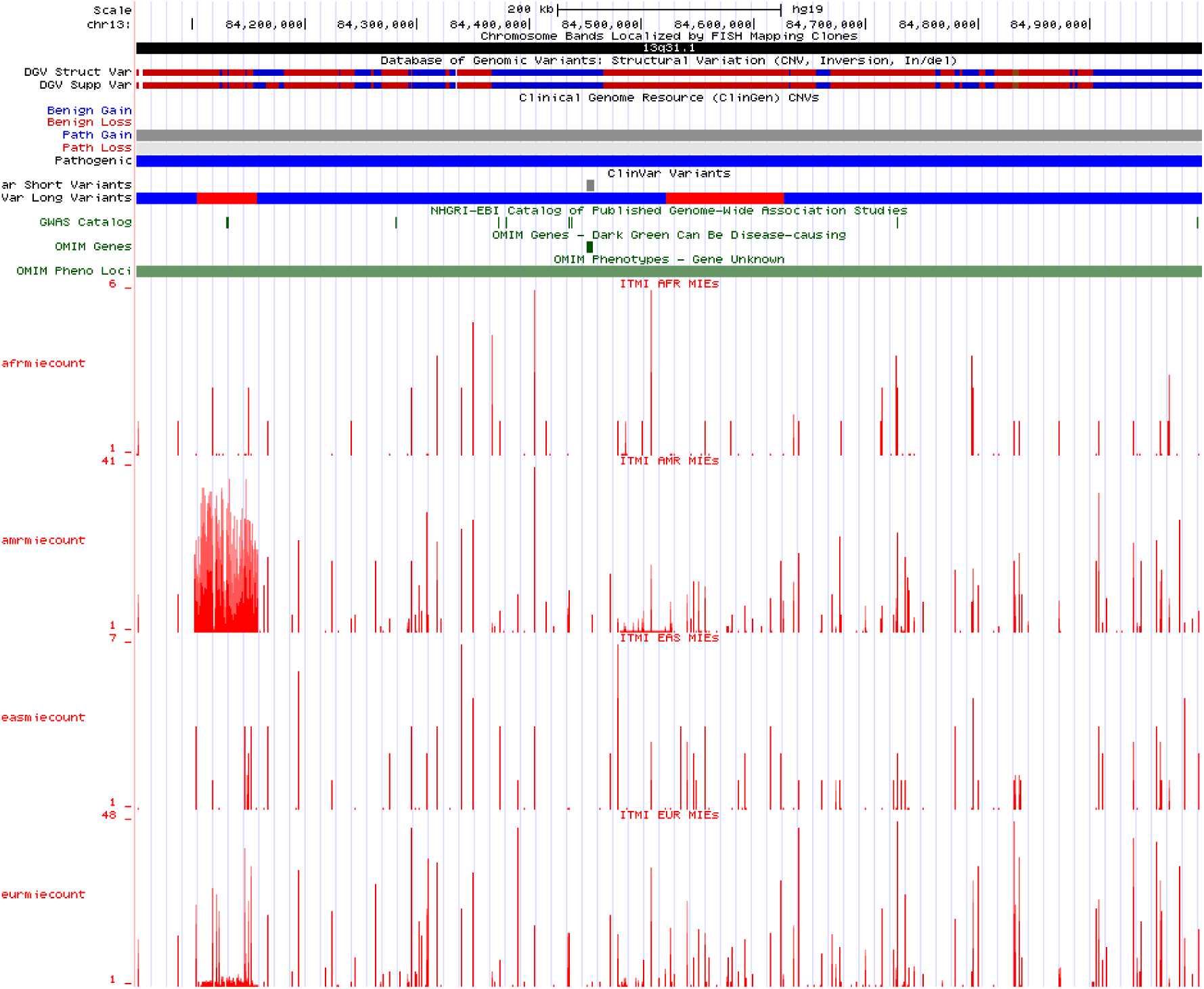
UCSC Genome Browser tracks displaying population-specific MIE profiles. Height of a bar represents number of trios with MIE at the position.

In order to determine if MIEs can be used for population stratification at a larger scale, we performed principal component analysis (PCA) on MIEs after removing calls with low quality or overlapping repeats. We excluded repetitive regions in order to ensure that the explained variance was not primarily due to population-specific repeats. We only retained MIEs that were found in fewer than 100 trios to exclude positions with systematic errors, and subsequently aggregated MIE counts in 1Mb window for each trio. The second, third and fourth principal components (PC2, PC3, PC4) could stratify the major populations as shown in Figure 4c. The first principal component (PC1) explained variability in number of MIE counts across trios (Supplementary Figure 7). The top 1Mb feature that contributed to PC2 showed different MIE distributions in AMR vs. EUR (Figure 4b) and corresponded to a heterozygous deletion in Chromosome 13 (detected with Manta, Supplementary Figure 8). Another 1Mb feature contributing to PC4 contained clustered MIEs (Supplementary Figures 9 and 10) and corresponded to a deletion in Chromosome 7 in EAS (detected with Control-FREEC).

**Figure 4c:**
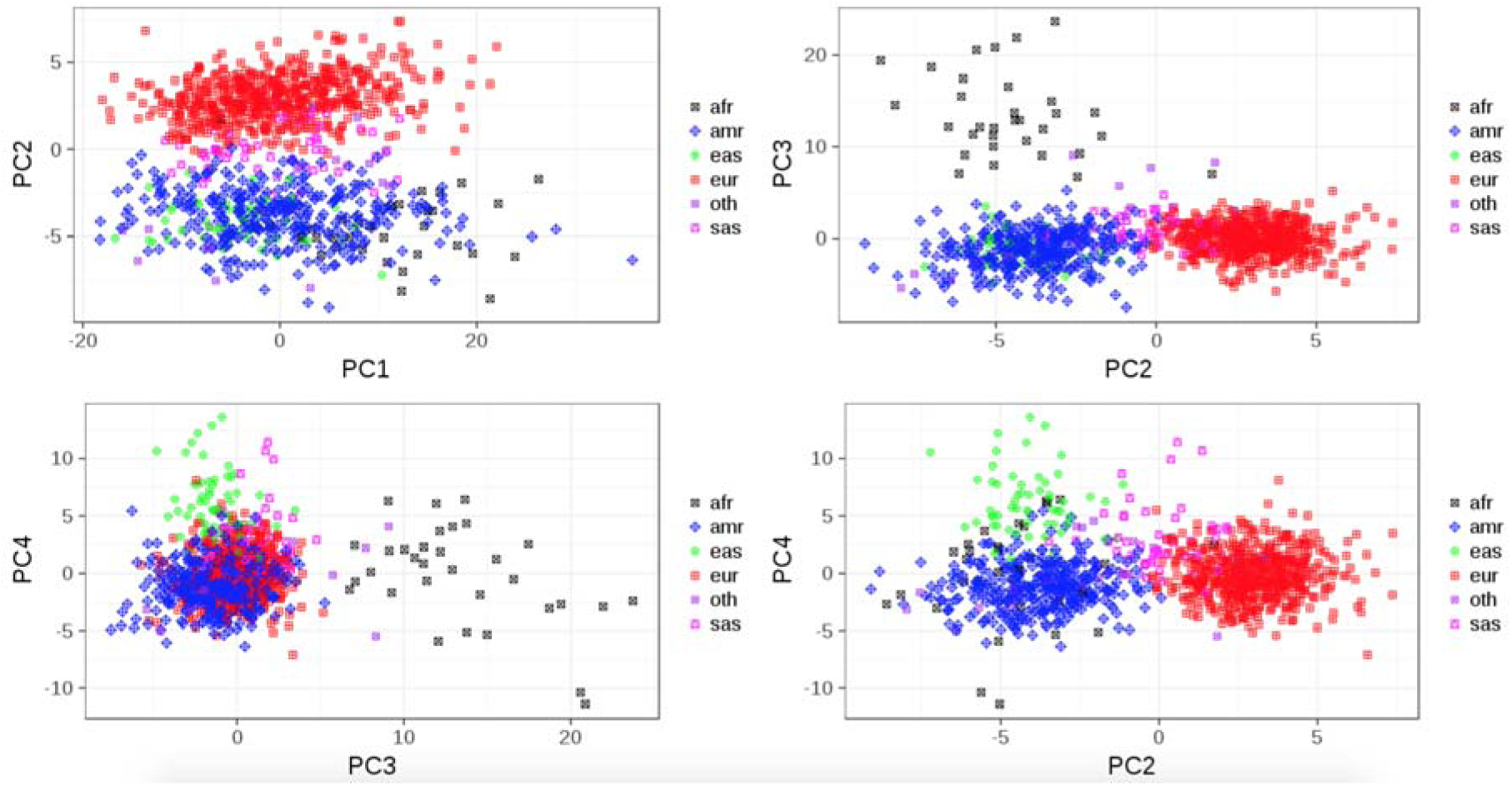
Principal component analysis (PCA) plots of principal components for Mendelian inheritance errors in trios. Points are colored by calculated ancestry. X-axis and Y-axis denote different principal components in each plot

**Figure 5:**
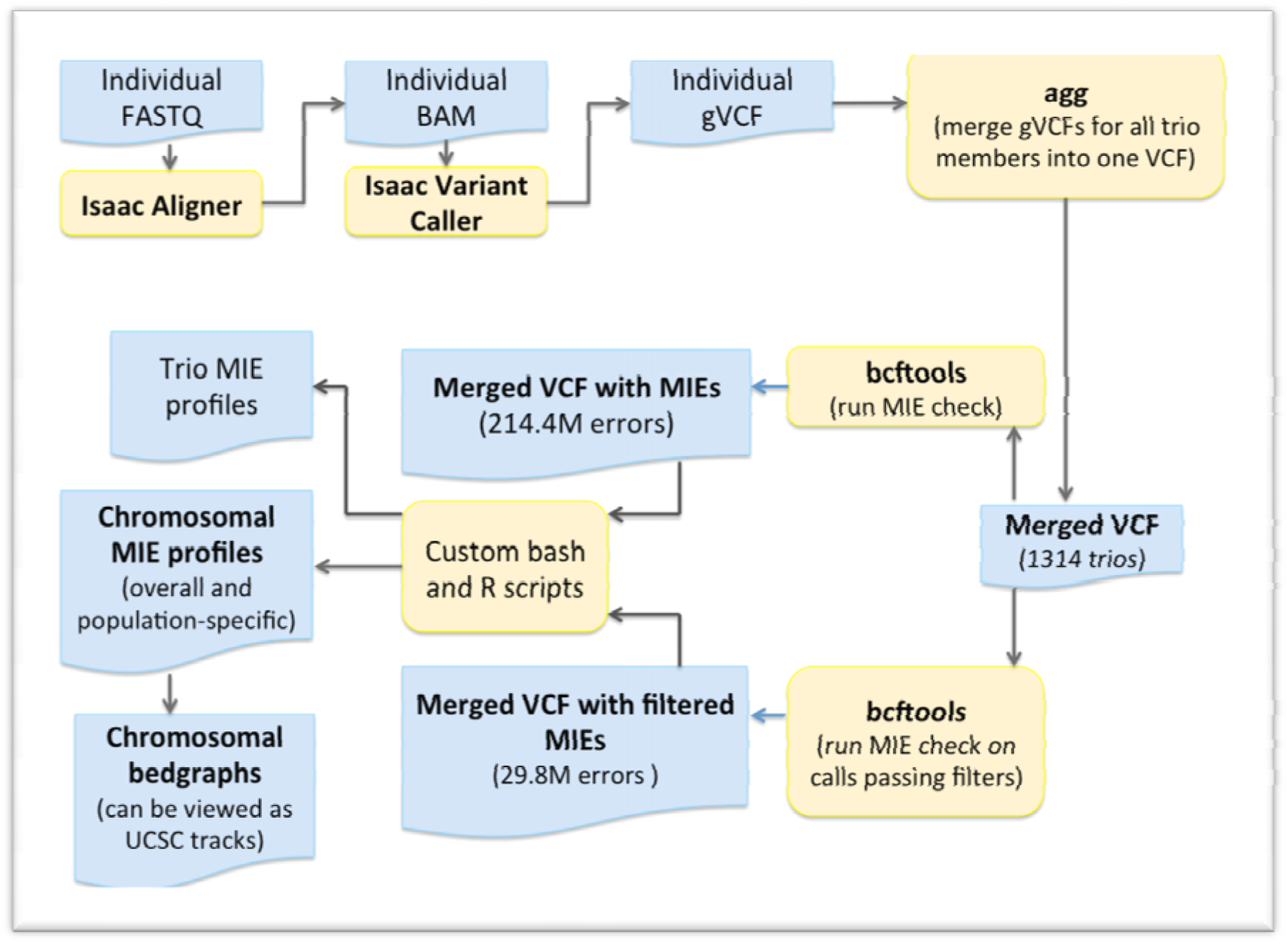
Analysis workflow

## Methods

### Sequencing data

The 1,314 complete family trios (father, mother and child) with validated pedigrees were obtained from an Institutional Review Board-approved childhood longitudinal WGS study. Informed consent was obtained for all subjects in the study. Trios were sent to Illumina Inc., (San Diego, CA) for WGS, assembly, and variant calling (pipeline version 2.01-2.0.3).

### Generating and annotating Mendelian Inheritance Errors

Individual gVCFs from all subjects were combined using agg (Illumina 2015). Resulting merged VCF was then processed to generate a list of MIEs in the autosomes using bcftools (version 1.3) (Danecek et al. 2011) mendelian plugin and custom shell scripts. No filters were applied initially to these MIE calls in order allow a comprehensive evaluation of quality metrics for the sites and calls bearing these errors. MIE-bearing loci were annotated with annovar (Wang et al. 2010). RepeatMasker (Smit et al. 1996) track from UCSC Table Browser (Kent et al. 2002) was used for annotating sites overlapping repeat elements.

### Computing distribution of quality metrics

Computations were performed using a combination of bcftools and custom R (version 3.4.0) scripts. R package ggplot2 (version 2.2.1) (https://github.com/tidvverse/ggplot2) was used for creating plots. Processing for all autosomes was run in parallel on our local cluster. Tab-delimited files containing quality metrics for all inconsistent trios were created for each autosome. These files were processed to obtain quality distributions across Mendelian-consistent and - inconsistent trio calls across MIE sites. Based on previously selected criteria (Bodian et al. 2016), we used cutoffs for GQ, DP and AB at 30, 10, and 0.25 respectively to determine the impact of filtering on MIE distribution.

### Overlap with NIST high confidence regions and 1000Genomes structural variation map

NIST high confidence regions for NA12878 (Zook et al. 2014) were obtained in the form of a BED file (ftp://ftp-trace.ncbi.nlm.nih.gov/giab/ftp/release/NA12878_HG001/NISTv3.2.2). An integrated map of structural variation calls in 1000Genomes Phase 3 data (Sudmant et al. 2015) was downloaded as a VCF from 1000Genomes FTP site (ftp://ftp.1000genomes.ebi.ac.uk/vol1/ftp/phase3/integrated_sv_map/ALL.wgs.integratedsvmapv2.20130502.svs.genotypes.vcf.gz). We then split each region in these files into windows of 100 bases using bedtools (Quinlan and Hall 2010) makewindows command in order to compare overall NIST, 1000Genomes SV, and ITMI MIE profiles. Peaks were then visualized by plotting cumulative counts over 100Kb bins.

### Detecting MIE enrichment and generating population-specific MIE tracks

In order to explore if concentration of MIEs in a region could indicate an overlapping deletion, we constructed MIE profiles for each trio across all autosomes. We calculated Standardized score for total number of MIE sites and number of unique MIE sites for each autosome per trio as follows:

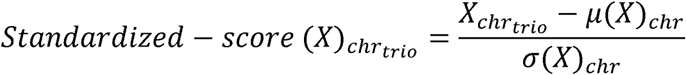

The samples were projected onto 1000Genomes (1000Genomes Project Consortium et al. 2015) principal components using the 17535 SNPs specified in AKT git repository using AKT (Arthur et al. 2017); the projections were then clustered into 5 clusters using PCs 1−3 using AKT. The clusters were defined by the 5 1000Genomes phase 3 super populations (namely, AFR, AMR, EAS, EUR, and SAS). Those samples with silhouette score >0.6 are assigned the ancestry that the cluster belongs to whereas others are assigned "OTHERS" as their ancestry. For population-specific analyses, we created groups to include all trios where both parents are assigned the same ancestry. This resulted in 545 EUR, 316 AMR, 55 EAS, 49 SAS, 29 AFR, and 22 OTHERS trios. These groupings were then used to generate population-specific bedGraphs (http://www.genome.ucsc.edu/FAO/FAOformat.html#format1.8) to represent MIE sites and associated error frequencies for each sub-population.

## Discussion

The utility of considering known familial relationships for error estimation and correction during detection of variants from sequencing data has been demonstrated before (Roach et al. 2010; Chen et al. 2013; Martin et al. 2014). In trio-based studies, MIEs are often used for estimating and alleviating sequencing error rates, and for optimizing filtering parameters. However, MIEs are often discarded as inconsistencies even before being studied for their cause, impact, and significance. We employed WGS data from 1,314 trios from diverse populations to understand and explore the characteristics and utility of MIEs.

We corroborated previous findings that MIEs are enriched in repeat elements and that quality distributions vary between Mendelian-consistent and –inconsistent calls (Patel et al. 2014; Pilipenko et al. 2014; Martin et al. 2014; Blackburn et al. 2014). Quality-based filtering has a greater impact on MIEs that occur across a higher proportion of trios compared to unique MIEs found in a single trio at a site. This observation is expected if we consider that infrequent inconsistencies could be due to incorrect genotype calls on real sequence variation and may not be errors attributed to problematic regions that present similar challenges across all samples. Previous studies involving MIEs have focused on developing filters to discard MIEs as sequencing errors or to extract a list of candidate de novo mutations. Our results indicate that MIE profiles and regions enriched for unique MIEs can be explored further as a tool for discovering putative deletions.

The repetitive nature of the human genome has been known to introduce mapping and alignment challenges (MacArthur et al. 2012; Treangen and Salzberg 2012). We observed highest MIE density in chromosome 19, which has been studied previously due to its high repeat density (Grimwood et al. 2004). We found that correlation between repeat and MIE densities across autosomes varies based on the type of repeat element where SINE density shows the strongest correlation with MIE density. Alu elements are primate-specific SINEs, which are ⍰300 nt in length, and propagate within a genome through retrotransposition. The abundance of Alu elements combined with their structure and presence of poly-A tail of variable length poses sequencing, mapping and variant calling challenges (Lander et al. 2001; Deininger 2011). Alu repeats are divided into three sub-families (AluJ, AluS, and AluY, from oldest to youngest) and together they contributed to ~24% MIEs post-filtering. Further work is required to understand if different MIE profiles are presented in the younger and polymorphic AluY sub-family vs. the older and more stable AluS and AluJ sub-families, and if these can be used to detect polymorphic Alu insertion or deletion with respect to the reference genome (Hormozdiari et al. 2011).

Using standardized scores, we identified trios that are outliers in terms of number of MIEs and/or unique MIEs for any given autosome. The comparison was performed at the chromosomal level as a first step to detect copy number losses. However, the approach could also be extended to detect deletions in smaller regions. To test the approach, we inspected the 22 outliers for each autosome and were able detect copy number losses that were called by Manta (or only by Control-FREEC). These results can be used to develop a validation approach for deletions called with alternative tools. Our results indicate that there is an opportunity for combining MIE data with coverage information to improve the detection of deletions. The finding is of significance especially since very few SV discovery tools offer genotyping as an option for calls derived from short read data (Marschall et al. 2013).

We also observed MIE peaks in regions that are enriched for deletions in 1000Genomes SV map. However, there are exceptions that warrant further exploration. Separating the MIE profiles by population enabled us to detect regions that have population-specific chromosomal anomalies. Researchers could use these MIE profiles in conjunction with NIST HCR and 1000Genomes SV map to better define callable regions in the genome.

Our analysis has limitations that require discussion. Further work is required to develop and optimize quality-based filters using MIE data for benchmarking. We adopted previously used filters for GQ, DP, and AB but did not refine the thresholds using MIEs. Detection of deletions with clustered MIEs has limitations in that the presence of a heterozygous deletion has to overlap with single nucleotide variants to lead to MIE calls in the region. Also, the approach has not been explored to detect insertions. A future goal is to assess the impact of joint trio-based variant calling on MIEs. We expect joint calling to reduce number of MIEs but aim to compare the proportion of calls overlapping deletions that are converted to missing, reported as hemizygous, or assigned low quality.

The proportion of Mendelian inconsistencies in a dataset is often used as a measure of sequencing and genotyping quality. However, our results indicate that caution needs to be exercised when selecting MIE calls for benchmarking, especially when a subset of the genome with a common copy number loss is being analyzed. Clustered MIEs, and MIEs with the signature of calls overlapping putative deletions (child and one parent are homozygous for opposite alleles) should be evaluated further to ensure these are not in reality hemizygous calls due to an overlapping deletion. Additionally, Mendelian consistent calls within a cluster of MIEs also need to be flagged and inspected before allele frequency and admixture calculation as these could be hemizygous calls that do not manifest as Mendelian inconsistencies but the genotypes could still be incorrect. Our observations highlight the need for a trio-aware variant caller that takes coverage differences into account and adds a flag to variants within a region that shows variation in coverage among trio members. Another option is first calling structural variations and then adjusting zygosity in regions overlapping deletions. However, this option can be computationally expensive when analyzing several thousand genomes. As a workaround, our future work comprises of using MIE bed files and annotating calls that are close to clustered MIEs. Researchers working with a small number of trios, especially trios from a non-European population, can include ITMI MIE tracks in their annotation pipeline to enable comparison of population-specific MIE rates in any given region, for selection of Mendelian inconsistencies for benchmarking, and annotating candidate de novo variations. In conclusion, Mendelian inconsistencies and other types of errors need to be understood and characterized better before being discarded as they could be representative of an underlying feature of the data.

## References

1000Genomes Project Consortium, Auton A, Brooks LD, Durbin RM, Garrison EP, Kang HM, Korbel JO, Marchini JL, McCarthy S, McVean GA, et al. 2015. A global reference for human genetic variation. Nature 526: 68–74.

Abecasis GR, Cherny SS, Cookson WO, Cardon LR. 2002. Merlin - rapid analysis of dense genetic maps using sparse gene flow trees. Nat Genet 30: 97–101.

Arthur R, Schulz-Trieglaff O, Cox AJ, O’Connell J. 2017. AKT: Ancestry and kinship toolkit. Bioinformatics 33: 142–144.

Bentley DR, Balasubramanian S, Swerdlow HP, Smith GP, Milton J, Brown CG, Hall KP, Evers DJ, Barnes CL, Bignell HR, et al. 2008. Accurate whole human genome sequencing using reversible terminator chemistry. Nature 456: 53–59.

Blackburn AN, Dean AK, Lehman DM. 2014. Imputation in families using a heuristic phasing approach. BMC Proc 8: S16.

Blue EM, Sun L, Tintle NL, Wijsman EM. 2014. Value of mendelian laws of segregation in families: Data quality control, imputation, and beyond. Genet Epidemiol 38.

Bodian DL, Klein E, Iyer RK, Wong WSW, Kothiyal P, Stauffer D, Huddleston KC, Gaither AD, Remsburg I, Khromykh A, et al. 2016. Utility of whole-genome sequencing for detection of newborn screening disorders in a population cohort of 1,696 neonates. Genet Med 18.

Boeva V, Popova T, Bleakley K, Chiche P, Cappo J, Schleiermacher G, Janoueix-Lerosey I, Delattre O, Barillot E. 2012. Control-FREEC: a tool for assessing copy number and allelic content using next-generation sequencing data. Bioinformatics 28: 423–425.

Chen W, Li B, Zeng Z, Sanna S, Sidore C, Busonero F, Kang HM, Li Y, Abecasis GR. 2013. Genotype calling and haplotyping in parent-offspring trios. Genome Res 23: 142–151.

Chen X, Schulz-Trieglaff O, Shaw R, Barnes B, Schlesinger F, Källberg M, Cox AJ, Kruglyak S, Saunders CT. 2016. Manta: Rapid detection of structural variants and indels for germline and cancer sequencing applications. Bioinformatics 32: 1220–1222.

Conrad DF, Andrews TD, Carter NP, Hurles ME, Pritchard JK. 2006. A high-resolution survey of deletion polymorphism in the human genome. Nat Genet 38: 75–81.

Conrad DF, Keebler JEM, DePristo MA, Lindsay SJ, Zhang Y, Casals F, Idaghdour Y, Hartl CL, Torroja C, Garimella K V, et al. 2011. Variation in genome-wide mutation rates within and between human families. Nat Genet 43: 712–4.

Danecek P, Auton A, Abecasis G, Albers CA, Banks E, DePristo MA, Handsaker RE, Lunter G, Marth GT, Sherry ST, et al. 2011. The variant call format and VCFtools. Bioinformatics 27: 2156–2158.

Deininger P. 2011. Alu elements: know the SINEs. Genome Biol 12: 236.

Dewey FE, Grove ME, Pan C, Goldstein BA, Bernstein JA, Chaib H, Merker JD, Goldfeder RL, Enns GM, David SP, et al. 2014. Clinical Interpretation and Implications of Whole-Genome Sequencing. JAMA 311: 1035.

Douglas J a, Boehnke M, Lange K. 2000. A multipoint method for detecting genotyping errors and mutations in sibling-pair linkage data. Am J Hum Genet 66: 1287–1297.

Epi4K Consortium E and E, Epilepsy Phenome/Genome Project, Allen AS, Berkovic SF, Cossette P, Delanty N, Dlugos D, Eichler EE, Epstein MP, Glauser T, et al. 2013. De novo mutations in epileptic encephalopathies. Nature 501: 217–21.

Goldmann JM, Seplyarskiy VB, Wong WSW, Vilboux T, Bodian L, Solomon BD, Veltman JA, Deeken JF, Gilissen C. 2017. Germline de novo mutation clusters arise during oocyte aging in genomic regions with increased double-strand break incidence. bioRxiv.

Goldmann JM, Wong WSW, Pinelli M, Farrah T, Bodian D, Stittrich AB, Glusman G, Vissers LELM, Hoischen A, Roach JC, et al. 2016. Parent-of-origin-specific signatures of de novo mutations. Nat Genet 48: 935–939.

Grimwood J, Gordon L a, Olsen a, Terry a, Schmutz J, Lamerdin J, Hellsten U, Goodstein D, Couronne O, Tran-Gyamfi M, et al. 2004. The DNA sequence and biology of human chromosome 19. Nature 428: 529–535.

Grover D, Mukerji M, Bhatnagar P, Kannan K, Brahmachari SK. 2004. Alu repeat analysis in the complete human genome: trends and variations with respect to genomic composition. Bioinformatics 20: 813–817.

Hormozdiari F, Alkan C, Ventura M, Hajirasouliha I, Malig M, Hach F, Yorukoglu D, Dao P, Bakhshi M, Sahinalp SC, et al. 2011. Alu repeat discovery and characterization within human genomes. Genome Res 21: 840–9.

Illumina. 2015. agg: A utility for aggregating Illumina-style GVCFs.

Jünemann S, Sedlazeck FJ, Prior K, Albersmeier A, John U, Kalinowski J, Mellmann A, Goesmann A, von Haeseler A, Stoye J, et al. 2013. Updating benchtop sequencing performance comparison. Nat Biotechnol 31: 294–6.

Kent WJ, Sugnet CW, Furey TS, Roskin KM, Pringle TH, Zahler AM, Haussler a. D. 2002. The Human Genome Browser at UCSC. Genome Res 12: 996–1006.

Kong A, Frigge ML, Masson G, Besenbacher S, Sulem P, Magnusson G, Gudjonsson SA, Sigurdsson A, Jonasdottir A, Jonasdottir A, et al. 2012. Rate of de novo mutations and the importance of father’s age to disease risk. Nature 488: 471–475.

Lander ES, Linton LM, Birren B, Nusbaum C, Zody MC, Baldwin J, Devon K, Dewar K, Doyle M, FitzHugh W, et al. 2001. Initial sequencing and analysis of the human genome. Nature 409: 860–921.

Loman NJ, Misra R V, Dallman TJ, Constantinidou C, Gharbia SE, Wain J, Pallen MJ. 2012. Performance comparison of benchtop high-throughput sequencing platforms. Nat Biotechnol 30: 434–9.

Lou DI, Hussmann JA, Mcbee RM, Acevedo A, Andino R, Press WH. 2013. High-throughput DNA sequencing errors are reduced by orders of magnitude using circle sequencing. Proc Natl Acad Sci 110: 19872–19877.

MacArthur DG, Balasubramanian S, Frankish A, Huang N, Morris J, Walter K, Jostins L, Habegger L, Pickrell JK, Montgomery SB, et al. 2012. A Systematic Survey of Loss-of-Function Variants in Human Protein-Coding Genes. Science (80-) 335: 823–828.

Manheimer KB, Patel N, Richter F, Gorham J, Tai AC, Homsy J, Boskovski MT, Parfenov M, Goldmuntz E, Chung WK, et al. 2017. Robust identification of deletions in exome and genome sequence data based on clustering of Mendelian errors. bioRxiv 209478.

Marschall T, Hajirasouliha I, Schönhuth A, Brudno M. 2013. MATE-CLEVER: Mendelian-inheritance-aware discovery and genotyping of midsize and long indels. Bioinformatics 29: 3143–3150.

Martin LJ, Pilipenko V, Kaufman KM, Cripe L, Kottyan LC, Keddache M, Dexheimer P, Weirauch MT, Benson DW. 2014. Whole exome sequencing for familial bicuspid aortic valve identifies putative variants. Circ Cardiovasc Genet 7: 677–683.

McCarthy SE, Gillis J, Kramer M, Lihm J, Yoon S, Berstein Y, Mistry M, Pavlidis P, Solomon R, Ghiban E, et al. 2014. De novo mutations in schizophrenia implicate chromatin remodeling and support a genetic overlap with autism and intellectual disability. Mol Psychiatry 19: 652–8.

Meacham F, Boffelli D, Dhahbi J, Martin DI, Singer M, Pachter L. 2011. Identification and correction of systematic error in high-throughput sequence data. BMC Bioinformatics 12: 451.

O’Connell JR, Weeks DE. 1998. PedCheck: A program for identification of genotype incompatibilities in linkage analysis. Am J Hum Genet 63: 259–266.

O’Rawe J, Jiang T, Sun G, Wu Y, Wang W, Hu J, Bodily P, Tian L, Hakonarson H, Johnson WE, et al. 2013. Low concordance of multiple variant-calling pipelines: practical implications for exome and genome sequencing. Genome Med 2013 53 40: 2426–2431.

Patel ZH, Kottyan LC, Lazaro S, Williams MS, Ledbetter DH, Tromp H, Rupert A, Kohram M, Wagner M, Husami A, et al. 2014. The struggle to find reliable results in exome sequencing data: filtering out Mendelian errors. Front Genet 5: 16.

Pilipenko V V, He H, Kurowski BG, Alexander ES, Zhang X, Ding L, Mersha TB, Kottyan L, Fardo DW, Martin LJ. 2014. Using Mendelian inheritance errors as quality control criteria in whole genome sequencing data set. BMC Proc 8: S21.

Quinlan AR, Hall IM. 2010. BEDTools: A flexible suite of utilities for comparing genomic features. Bioinformatics 26: 841–842.

Rauch A, Wieczorek D, Graf E, Wieland T, Endele S, Schwarzmayr T, Albrecht B, Bartholdi D, Beygo J, Di Donato N, et al. 2012. Range of genetic mutations associated with severe non-syndromic sporadic intellectual disability: an exome sequencing study. Lancet 380: 1674–1682.

Reumers J, De Rijk P, Zhao H, Liekens A, Smeets D, Cleary J, Van Loo P, Van Den Bossche M, Catthoor K, Sabbe B, et al. 2011. Optimized filtering reduces the error rate in detecting genomic variants by short-read sequencing. Nat Biotechnol 30: 61–68.

Roach JC, Glusman G, Smit AFA, Huff CD, Hubley R, Shannon PT, Rowen L, Pant KP, Goodman N, Bamshad M, et al. 2010. Analysis of Genetic Inheritance in a Family Quartet by Whole-Genome Sequencing. Science (80-) 328: 636–639.

Ronemus M, Iossifov I, Levy D, Wigler M. 2014. The role of de novo mutations in the genetics of autism spectrum disorders. Nat Rev Genet 15: 133–141.

Ross MG, Russ C, Costello M, Hollinger A, Lennon NJ, Hegarty R, Nusbaum C, Jaffe DB, Shea T, Sykes S, et al. 2013. Characterizing and measuring bias in sequence data. Genome Biol 2013 145 108: 1513–1518.

Smit A, Hubley R, Green P. 1996. RepeatMasker Open-3.0. RepeatMasker Open-30 www.repeatmasker.org.

Sobel E, Papp JC, Lange K. 2002. Detection and integration of genotyping errors in statistical genetics. Am J Hum Genet 70: 496–508.

Sudmant PH, Rausch T, Gardner EJ, Handsaker RE, Abyzov A, Huddleston J, Zhang Y, Ye K, Jun G, Hsi-Yang Fritz M, et al. 2015. An integrated map of structural variation in 2,504 human genomes. Nature 526: 75–81.

Treangen TJ, Salzberg SL. 2012. Repetitive DNA and next-generation sequencing: Computational challenges and solutions. Nat Rev Genet 13: 36–46.

Wang K, Li M, Hakonarson H. 2010. ANNOVAR: Functional annotation of genetic variants from high-throughput sequencing data. Nucleic Acids Res 38.

Weir D. 2012. Quality Control Report for Genotypic Data.

Wong WSW, Solomon BD, Bodian DL, Kothiyal P, Eley G, Huddleston KC, Baker R, Thach DC, Iyer RK, Vockley JG, et al. 2016. New observations on maternal age effect on germline de novo mutations. Nat Commun 7.

Yuen RK, Merico D, Cao H, Pellecchia G, Alipanahi B, Thiruvahindrapuram B, Tong X, Sun Y, Cao D, Zhang T, et al. 2016. Genome-wide characteristics of de novo mutations in autism. npj Genomic Med 1: 16027.

Zook JM, Chapman B, Wang J, Mittelman D, Hofmann O, Hide W, Salit M. 2014. Integrating human sequence data sets provides a resource of benchmark SNP and indel genotype calls. Nat Biotechnol 32: 246–251.

